# Molecular Deconstruction of the Medullary Raphe Magnus in the rat: Transcriptional Responses to Repeated Seizures

**DOI:** 10.1101/2025.09.22.677744

**Authors:** Paloma Bittencourt-Silva, Ibrahim Vazirabad, Melissa Eilbes, Luke Zangl, Wasif Osmani, Matthew R. Hodges

**Author notes:** Correspondence: Matthew R. Hodges, PhD, B-5205, Dept. of Physiology, Medical College of Wisconsin, Milwaukee, WI 53226. Authors contributed equally to this work (Paloma Bittencourt-Silva, Ibrahim Vazirabad). This work was funded by National Institutes of Health R01HL122358.

## Abstract

Sudden Unexpected Death in Epilepsy (SUDEP) is a leading cause of death in patients with epilepsy and is thought to result from dysfunctional and/or failure of cardiorespiratory control systems. Post-mortem brainstem tissue analyses in human SUDEP cases point to reductions in markers of the brainstem serotonin (5-HT) system, which is known as a brainstem center that provides excitatory neuromodulation. We have previously shown in a knockout rat model (SS*^kcnj16-/-^* rats) that repeated seizures led to progressively greater ventilatory inhibition in the post-seizure period, seizure-associated mortality, and reduced brainstem 5-HT and tryptophan hydroxylase (Tph) particularly within the Raphe Magnus (RMg). Here, we account for the cellular constituency and local transcriptional responses to repeated seizures in male SS*^kcnj16-/-^* rats that experienced daily seizures for 3, 5, 7, or 10 consecutive days using single nuclear RNA sequencing (snRNA seq) from brainstem tissue biopsies including the RMg (-12.12 mm to -10.30 mm caudal to Bregma) two hours post-seizure. Unbiased cluster analysis identified 18 cell major clusters that were by identified by the expression of known gene markers, with most cells being oligodendrocytes. However, local RMg neurons showed the greatest numbers of differentially expressed genes with seizures compared to all other cell types. Further re-clustering of neuronal cell types yielded 14 distinct RMg neuron subpopulations, including 5 types of GABAergic neurons, 2 glutamatergic clusters, and 2 groups of 5-HT neurons which all had unique expression profiles. Nearly all DEGs across neuronal subtypes were increased following seizures, and a large fraction of which were common across seizure days and across neuron type suggesting uniformity in cellular response to seizures in this region. These studies provide foundational information regarding the cellular constituency of the RMg region in the rat, and altered neuronal function following repeated seizures in the absence of changes in other cell types in this key region of cardiorespiratory control.

## Introduction

Millions of people suffer from spontaneous, recurrent seizures in epilepsy and other seizure disorders. An estimated 30% of patients with epilepsy are not responsive to anti-seizure medications (ASMs)^1^, and it is those patients that are at a particularly high risk of Sudden Unexpected Death in Epilepsy (SUDEP)^2-4^. Retrospective pathophysiologic data analyses of documented SUDEP events in humans revealed a common sequence of events leading to a SUDEP event, which includes a generalized tonic clonic seizure (GTCS) followed by breathing and heart rate suppression preceding terminal apnea and followed by terminal asystole^5^. This landmark study provided key evidence of seizure-induced dysfunction in vital systems such as cardiorespiratory control, which is regulated by overlapping brainstem neural networks that may be disrupted in epilepsy by repeated seizures. While a mechanistic understanding of SUDEP remain a focus of investigation, accumulating data from post-mortem tissue studies from human SUDEP cases showed specific abnormalities within brainstem regions thought to be responsible for generating breathing rhythm (the pre-Bötzinger Complex; preBötC) or that contain neurons that synthesize and release 5-Hydroxytryptamine (5-HT; serotonin) in the ventrolateral medulla^6^. Thus, defining the effects of repeated seizures on specific elements of these neural circuits may therefore provide a greater mechanistic understanding of human SUDEP.

We have developed a rat model to study the effects of repeated seizures on cardiorespiratory functions. Knockout of *Kcnj16*, the gene that encodes the inwardly rectifying potassium ion channel subunit Kir5.1, in Dahl Salt-sensitive inbred rats (SS*^kcnj16-/-^* rats) led to several physiologic^7,8^ and neurological phenotypes^9^, including a sensitivity to sound-induced seizures^10^. Eliciting one seizure/day for up to 10 days in SS*^kcnj16-/-^* rats led to a progressively greater cardiorespiratory dysfunction after each seizure^11^, and in some cases led to seizure-induced deaths (SzIDs) particularly when seizures were elicited in the early morning or early evening^12^. Daily seizures in SS*^kcnj16-/-^* rats also resulted in time-dependent increases in neuroinflammatory molecules (cytokines and chemokines) and markers of microglial activation (increased IBA1 and altered morphology) within the critically important pre-BötC^13^. This region and its function in respiratory rhythmogenesis relies on excitatory and inhibitory neuromodulatory inputs, including from serotonergic raphe neurons which release of 5-HT, substance P and thyrotropin-releasing hormone (TRH) within the preBötC and other brainstem nuclei^14-16^. Similar to that shown in human SUDEP cases, tissue assessments of 5-HT and tryptophan hydroxylase (TPH) in SS*^kcnj16-/-^* rats exposed to daily seizures showed large deficits in TPH-immunoreactivity within Raphe Magnus (RMg) and reductions in 5-HT-immunoreactivity in the pre-BötC, nucleus of the solitary tract (nTS) and nucleus ambiguous (NA)^11^. While repeated seizures in this model caused wide-spread suppression of the brainstem raphe 5-HT system, this effect is likely not due to a neuroinflammatory process^13^. Therefore, we hypothesize that repeated seizures downregulate raphe TPH and brainstem 5-HT production through a local RMg mechanism. However, there are few data that provide a comprehensive local cellular constituency of the RMg. Furthermore, the transcriptional response of RMg neurons to repeated seizures may provide insights into potential neuronal dysfunction generated by repeated seizures in the RMg, which may lead to an increased understanding of the potential role of 5-HT deficiency in SUDEP. Here we performed single nuclear RNA sequencing (snRNAseq) from brainstem biopsies containing the RMg collected from male SS*^kcnj16-/-^* rats after repeated seizures for a complete accounting of the cellular constituency of the RMg of the rat medulla and measured differential gene expression patterns within neurons to gain additional insights into cellular responses to repeated seizures.

## Methods

### Animals

Male SS^Kcnj16-/-^ rats were obtained from colonies maintained at the Medical College of Wisconsin. Rats were housed in controlled environmental conditions under a 12-hour-light/dark cycle with food (Low K+; Dyets Inc., D113755) and water ad libitum and studied at 10 weeks old. All procedures and protocols were reviewed and approved by the Institutional Animal Care and Use Committee of the Medical College of Wisconsin.

### Seizures

For seizure induction, animals were placed in a plexiglass chamber, left for 20 minutes to acclimate and then were acoustically stimulated for 2 minutes. The acoustic stimulus of 10 kHz audio frequency at 100 dB amplitude was produced by a function generator (GW Instek model GFG-8020H) and delivered through a 50 Ω speaker (Visaton model k50wp) positioned about 5 inches above the rat. Seizure behaviors were video recorded, and seizure severity was scored based on a modified Racine scale optimized for the specific progression of behaviors consistently observed in these animals (Mannis et al 2023). Seizure induction was repeated once daily for either 3, 5, 7 or 10 days for each rat.

### Tissue collection

60-90 minutes after the last acoustic seizure, the rats were deeply anesthetized with isoflurane (5% in air) and the brain was collected, flash frozen and stored (-80°C). Four consecutive thick sections (∼0.5 mm each; 2 mm total) were generated while frozen (maintained at -20°C) and tissue biopsies/punches (2 mm diameter) from the raphe magnus were collected from all sections. This region spanned approximately -12.12 mm to -10.30 mm caudal to Bregma based on the Paxinos and Watson rat brain atlas^17^ and corresponds to a caudal-rostral range of the medulla from the Bötzinger complex to the rostral aspects of the facial motor nucleus. RMg tissue samples from SS*^kcnj16-/-^* rats exposed to zero (naïve; n=3), 3 days (n=3), 5 days (n=3), 7 days (n=4), or 10 days (n=3) of audiogenic seizures yielded similar numbers of nuclei for transcriptomic assessment. Cell nuclei were isolated from flash-frozen rat brain tissue samples and constructed 3’ single-cell gene expression libraries (Next GEM v3.1) using the 10x Genomics Chromium system. The libraries were sequenced with ∼200 million PE150 reads per sample on Illumina NovaSeq. The sequencing reads were analyzed with the *Rattus norvegicus* reference genome mRatBN7.2.106 from Ensembl using Cell Ranger v7.2.0. Introns were included in the analysis.

The resulting single nucleus transcript data were analyzed by Seurat v.5.1.0 in R v4.4.0. Quality control was performed by excluding nuclei with more than 5% mitochondrial content, less than 500 or more than 6,000 detected genes, and less than 1,000 or more than 20,000 detected RNA molecules. The scTransform function^18^ was used to normalize, scale, and find variable features for the nuclei. The initial 30 principal components were used for principal component analysis (PCA) and auto-clustering analysis. To mitigate any batch effects and other technical biases, the CCAIntegration function in Seurat was used to integrate all samples together for joint clustering. Clusters were identified for each nucleus at a resolution of 0.3. Results were visualized by generation of t-SNE plots using the R package scCustomize^19^. Differential expression within putative neuronal types between various seizure durations and naïve condition was generated by Wilcoxon test in the R presto package (criteria |Log2FC| > 0.5, FDR < 0.05).

## Results

The SS*^kcnj16-/-^* rats used in this study consistently responded to 2 min sound exposures with seizures of varying severity quantified using a Modified Racine scale as previously reported^10,11^. Seizure severity scores were not different across the 10-day seizure protocol (**Suppl. Fig. 1**; One-way ANOVA; P=0.41) and the mean seizure severity score within each rat group was within the range of scores shown previously in this model^10,11^. Similarly, changes in weight (grams; g) among rats across different seizure-exposed groups was not different (**Suppl. Fig. 1**; One-way ANOVA; P=0.42), suggesting rats were in good health at the time of tissue collection, although three SS*^kcnj16-/-^*rats died spontaneously prior to their designated tissue collection date and were not used in further analyses (3/16 total mortality) as was expected in this seizure model^10,13^.

### Medullary Raphe Magnus (RMg) cell types

Unbiased cluster analysis (PC=30, resolution=0.3) of the aggregate (all tissues and conditions) allowed for the segregation of RMg cell nuclei into 18 cell clusters (0-17; **Table 1**) based on differential expression of genes unique to each cluster. Cell cluster identity mapping was accomplished using a three-step process: 1) individual genes from the 20 most differentially expressed genes within that cluster and associated cell types in PanglaoDB, 2) a combination of 2 genes from the 10 most differentially expressed genes within each cluster and their associated cell type in PanglaoDB, and 3) a cell type analysis using the top 50 differentially expressed genes using the Allen Brain Atlas (**Table 1**). Steps 1 and 2 yielded a provisional cell type call using cell confidence scores, which agreed with cell type calls from the Allen Brain Atlas for all but one cluster (Cluster 4) which had neuronal gene expression features but low cell confidence scores and was ultimately determined to be more representative of a population of oligodendrocyte precursor cells/oligodendrocytes (OPC-oligo; **Table 1**). Final cell types were then identified using a t-SNE plot of the aggregate data to visualize clusters (**Fig. 1A**) and determined to be represented within all conditions (naïve and 3, 5, 7, and 10 days of seizures; **Fig. 1B**). The 18 clusters of identified cell types included 5 subclasses of oligodendrocytes (Oligo1-5), 6 subclasses of neurons (Neurons1, 3-7), oligodendrocyte precursors (OPCs) and a cell cluster that shared OPC and oligodendrocyte transcriptional properties (OPC-Oligo) along with discrete populations of astrocytes, microglia, endothelial cells (EC) fibroblasts and a final cluster with shared fibroblast and EC gene expression (Fibro_EC). Further confirmation of cell identities of all clusters was verified by the expression of known gene markers across all clusters (**Fig. 2**), which included *Mag* and *Mog* (Oligodendrocytes), *Aqp4* and S*lc1a2* (astrocytes), *Cntnap2* and *Nrxn1* (neurons), *Cx3cr1* and *Tgfbr1* (microglia), *Flt1* and *Ptprb* (endothelial cells), and *Bicc1* and *Vim* (fibroblasts). These combined analyses allowed for a determination of the complete cellular milieu present within the medullary RMg in the rat.

**Table 1.**
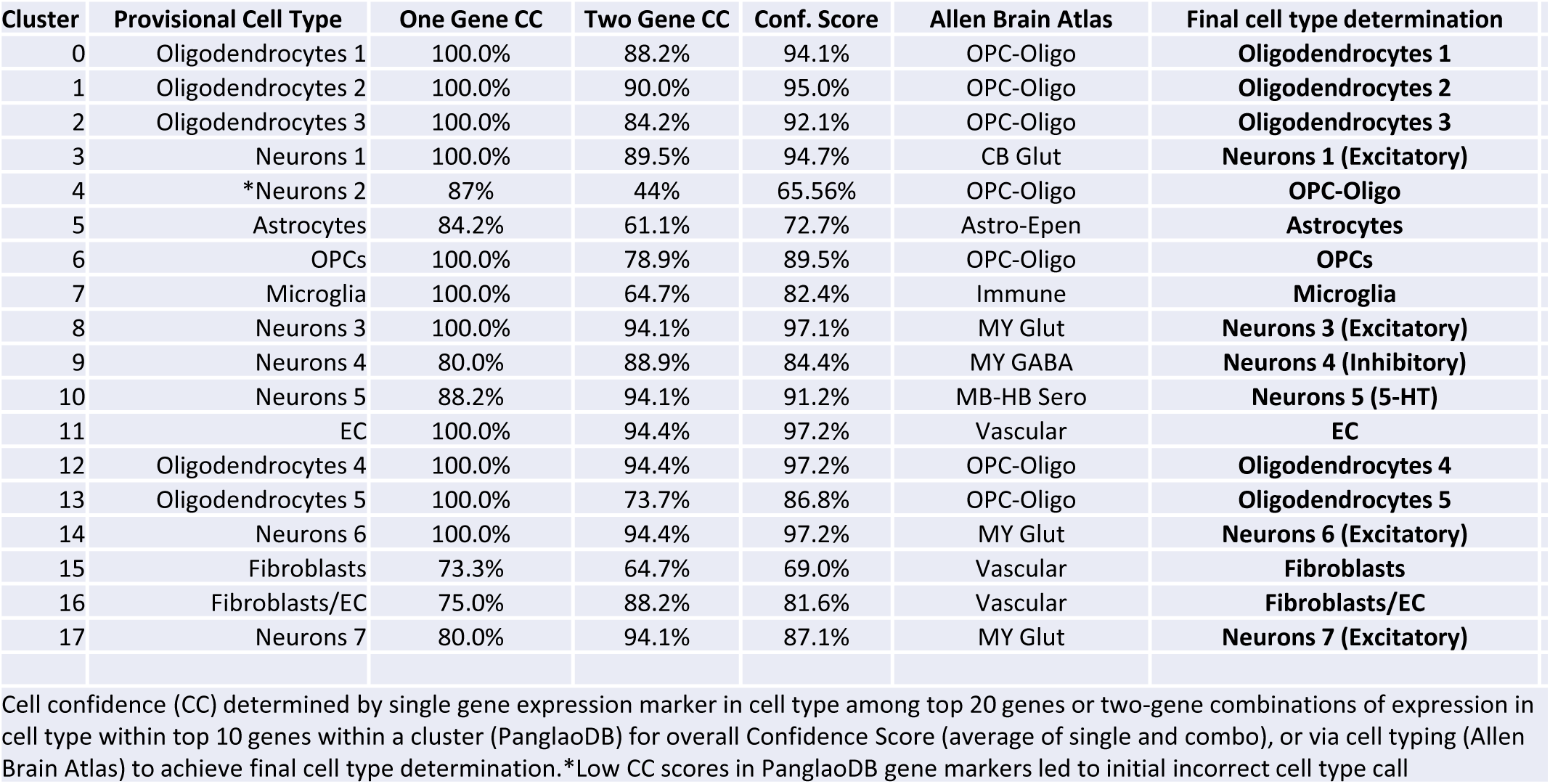

**Figure 1.**
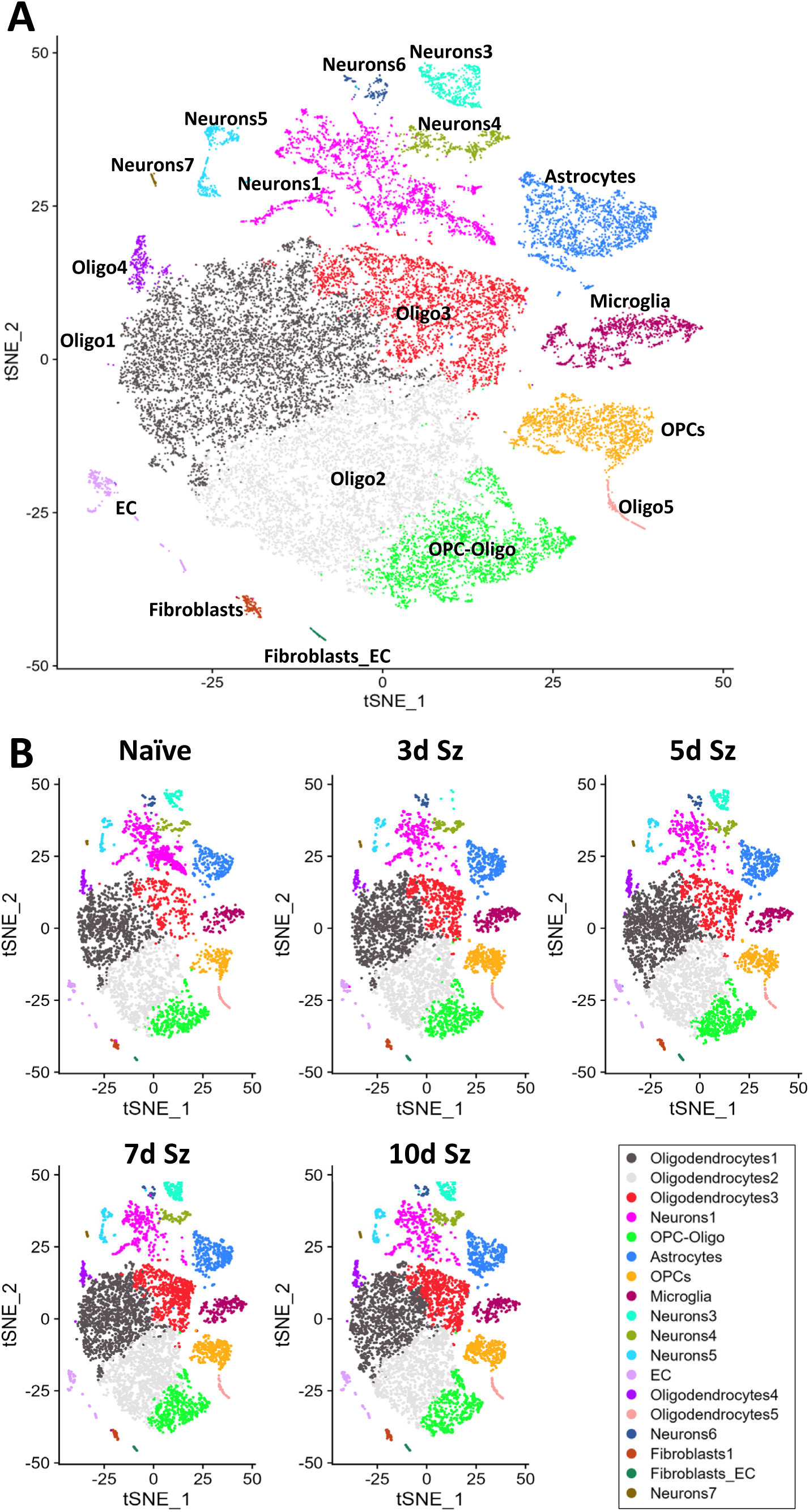
Unbiased cell clustering and cell types across condition. **A**) t-SNE plot of single-nucleus cell-types based on unbiased clustering: Neurons (1, 3-7), Oligodendrocytes (Oligo1-5), mixed Oligodendrocyte-Oligodendrocyte progenitor cells (OPC-Oligo), Oligodendrocyte progenitor cells (OPCs), Microglia, Astrocytes, Endothelial cells (EC), mixed Endothelial Cells-Fibroblasts (Fibroblasts_EC), and Fibroblasts. **B**) tSNE plots split by condition show that all clusters persist regardless of seizure duration.

**Figure 2.**
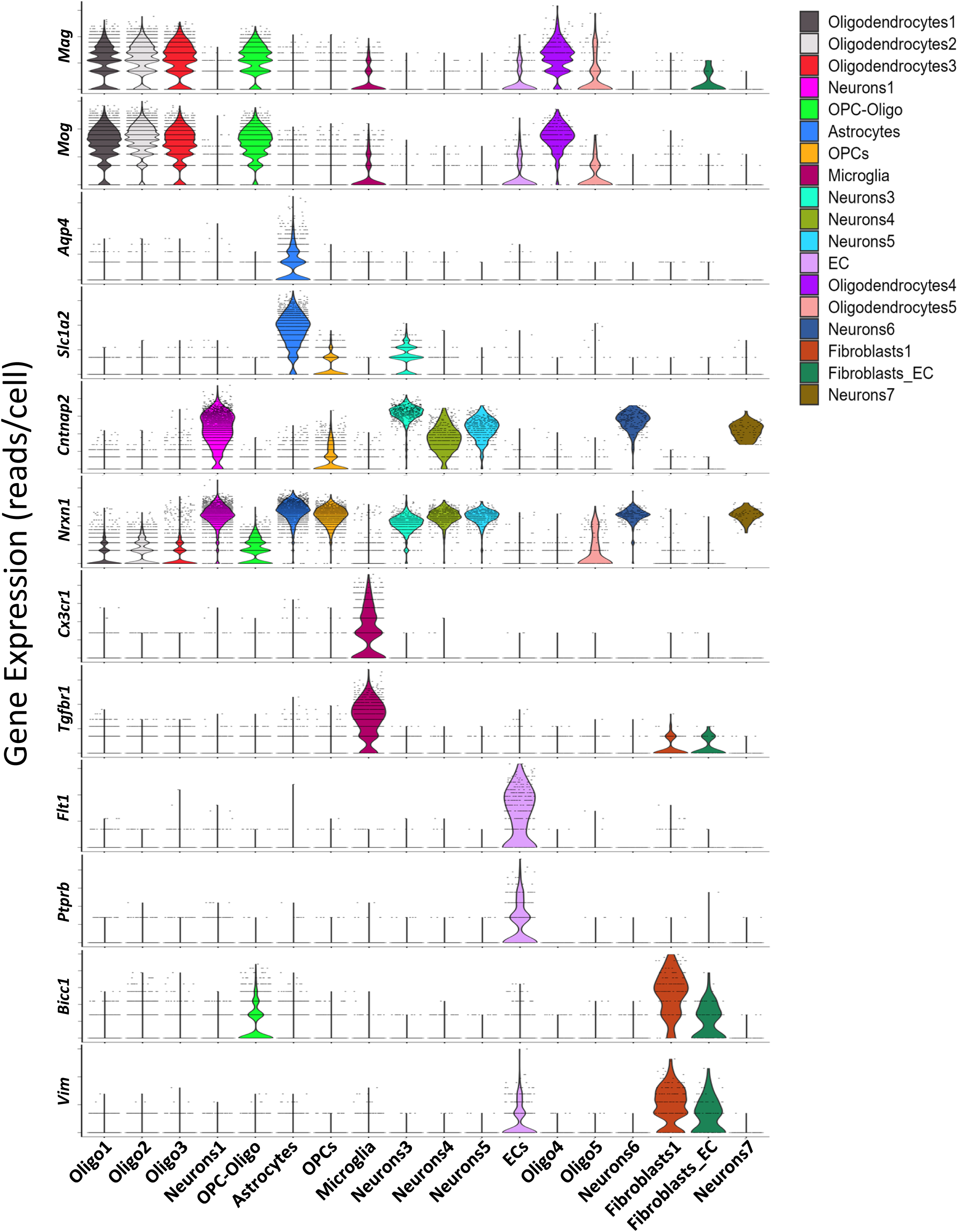
Cell type markers across all clusters. Expression violin plot of selected marker genes vs putative cell-type annotations.

### Major neuronal groups within the RMg

Further interrogation of additional genes expressed within the six major neuronal clusters in the RMg allowed for additional classification. The largest neuron subclass (<u>Neurons1</u>) showed modest expression of glutamic acid decarboxylase 1 (*Gad1*), 5-HT receptor 2C (*Htr2c*), gamma aminobutyric acid (GABA) A receptor subunit alpha 1 (*Gabra1*) and TREK-2 (*Kcnk10*) with little or no expression of glutamatergic neuron markers VGLUT2 (*Slc17a6*) or VGLUT3 (*Slc17a8*; **Fig. 3**) suggesting these were inhibitory neurons. <u>Neurons3</u> uniquely expressed VGLUT2 (*Slc17a6*) and corticotropin-releasing hormone (*Crh*), whereas <u>Neurons4</u> expressed both *Gad1* and *Gad2* along with *Htr2C* and *Gabra1* consistent with excitatory glutamatergic and GABAergic inhibitory neurons, respectively (**Fig. 3**). <u>Neurons5</u> uniquely expressed the serotonin cell markers tryptophan hydroxylase 2 (*Tph2*), dopa decarboxylase (*Ddc*) and SERT (*Slc6a4*) in addition to VGLUT3, Gad1, 5-HT2C receptor and TREK-2 (**Fig. 3**) among other genes such as tachykinin 1 (substance P; *Tac1*) and thyrotropin-releasing hormone (*Trh*; data not shown). <u>Neurons6</u> showed gene expression patterns similar to Neurons1, with modest levels of *Gad1*, *Gad2*, *Htr2C*, *Gabra1* and *Kcnk10*. Finally, <u>Neurons7</u>, which represented the fewest number of RMg neuronal nuclei, uniquely expressed peptide YY (*Pyy*) in addition to measurable levels of *Htr2C* (**Fig. 3**).

Separating these 6 major neuronal clusters from all other non-neuronal cells allowed for unbiased reclustering of neuronal subpopulations within the RMg to gain further insights into the neuron types within this region of the brainstem. Several iterations of altering dimensionality (resolution settings) yielded ∼14 unique neuronal clusters (NCs; NC_0 to NC_13; **Fig. 4**). As expected, each NC expressed common neuronal marker genes (*Nxrn1* and *Map2*) and most NCs could be classified as GABAergic (*Gad1/2*) or glutamatergic (*Slc17a6-8*; **Fig. 5**). With one exception (NC_0), each RMg neuron subtype expressed unique combinations of genes within each NC. Unlike other NCs, NC_0 was the largest group of RMg neurons but had no obvious unique transcriptomic identifiers that distinguished it from other NCs (**Fig. 5**), suggesting that NC_0 was a collection of mixed neuron types. This was confirmed by secondary cell typing analysis in the Allen Brain Atlas which showed NC_0 contained both glutamatergic and GABAergic neurons (data not shown). The second-largest neuron type (NC_1) was glutamatergic (*Slc17a7*) and co-expressed the distinct marker genes *Gabra6*, *Reln*, *Slc4a4*, *Adcy1*, *Mical2* and *Ndst3*, along with high levels of *Camk2d* (**Fig. 5**). NC_2 was also glutamatergic (*Slc17a6*) and uniquely co-expressed *Crh*, *Gpr88*, *Kcnh7* and *Nav2*. In addition to these two glutamatergic neuronal subtypes, there were 5 GABAergic NCs (*Gad1* and/or *Gad2*) with unique expression profiles that differentiated these subclasses. NC_3 co-expressed *Dcc*, *Erbb4*, and *Fbxl7*. NC_5 co-expressed *Adarb2*, *Camk2d*, and *Ghr*. NC_6 co-expressed *Grm3*, *Plp1*, and *Map7*. NC_7 co-expressed *Kcnd3*, *Sptlc3*, and *Nwd2*. NC_8 co-expressed *Adarb2*, *Rora*, *Phactr2*, and *Grm1* (**Fig. 5**).

**Figure 3.**
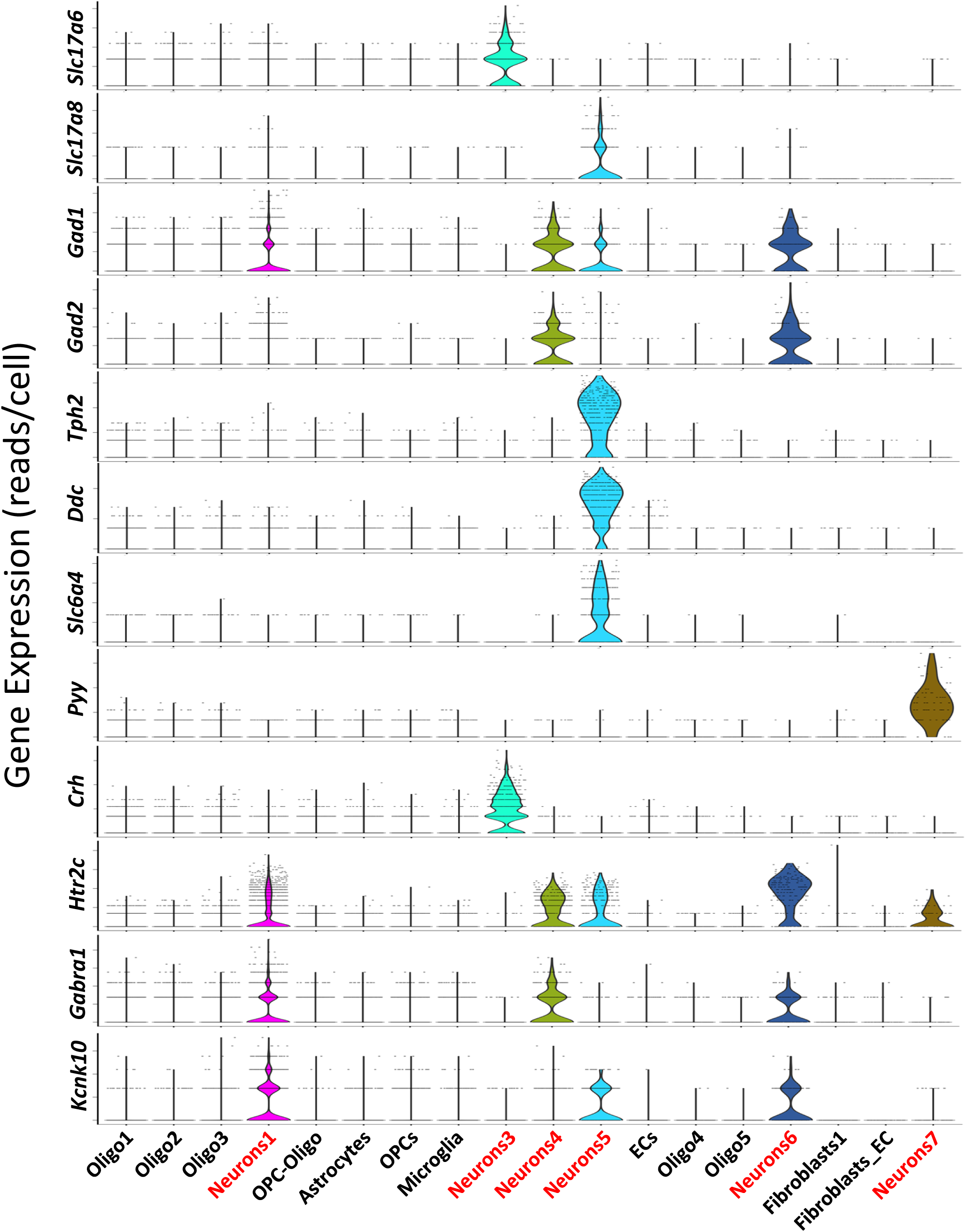
Neuronal subtype-specific markers. Expression violin plot of neuronal marker genes enriched in selected cell-type clusters. Note the expression of excitatory (Slc17a6/7) or inhibitory (Gad1/2) gene markers were restricted to neuronal clusters 3+5 or 1, 4, 5 + 6, respectively. Note also that serotonergic cell markers (Tph2/Ddc and Slc6a4) were restricted to neuronal cluster 5.

**Figure 4.**
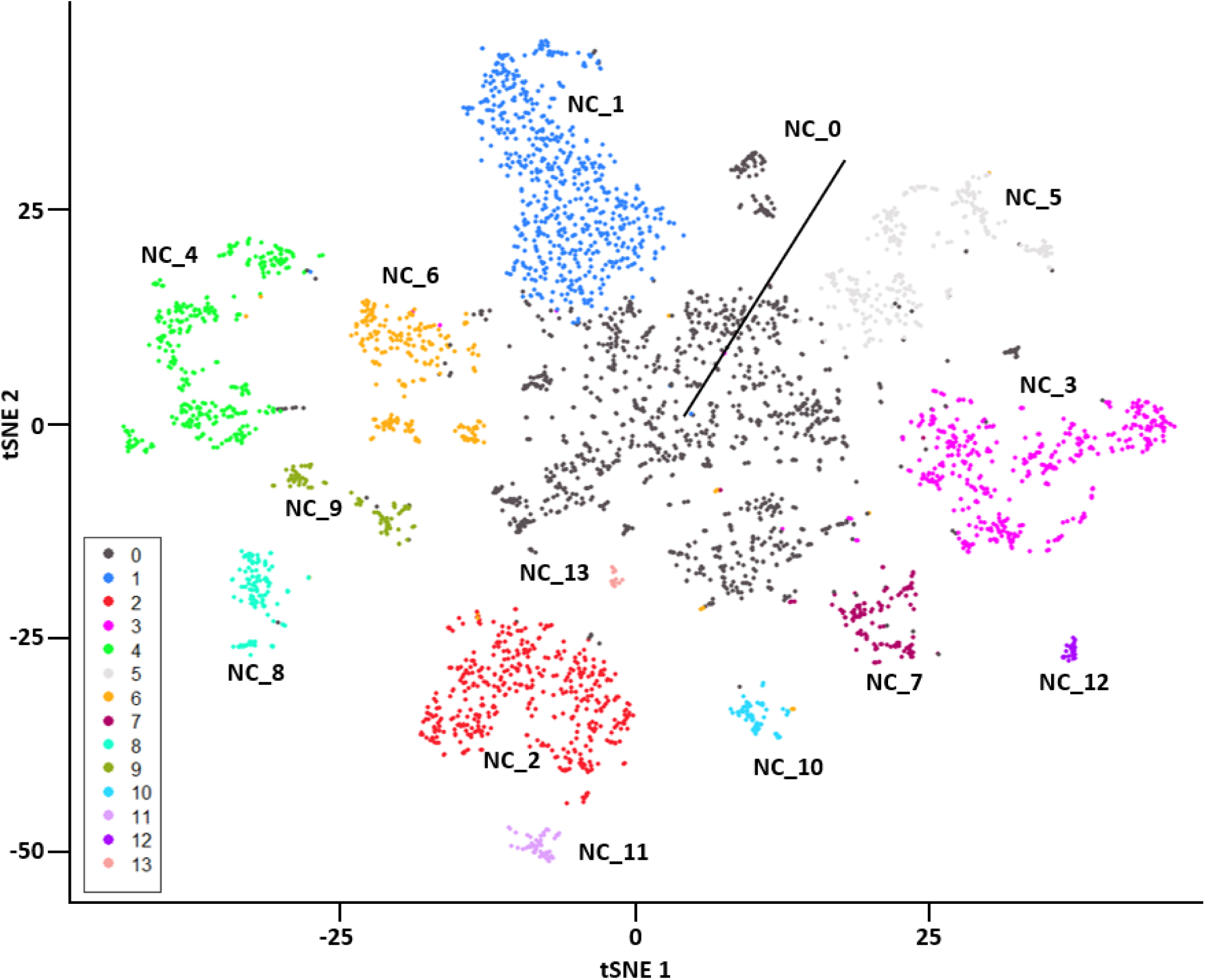
Unbiased neuronal re-clustering (aggregate). t-SNE plot of putative neuronal cell-types. Re-clustering provided fourteen distinct neuronal subpopulations.

**Figure 5.**
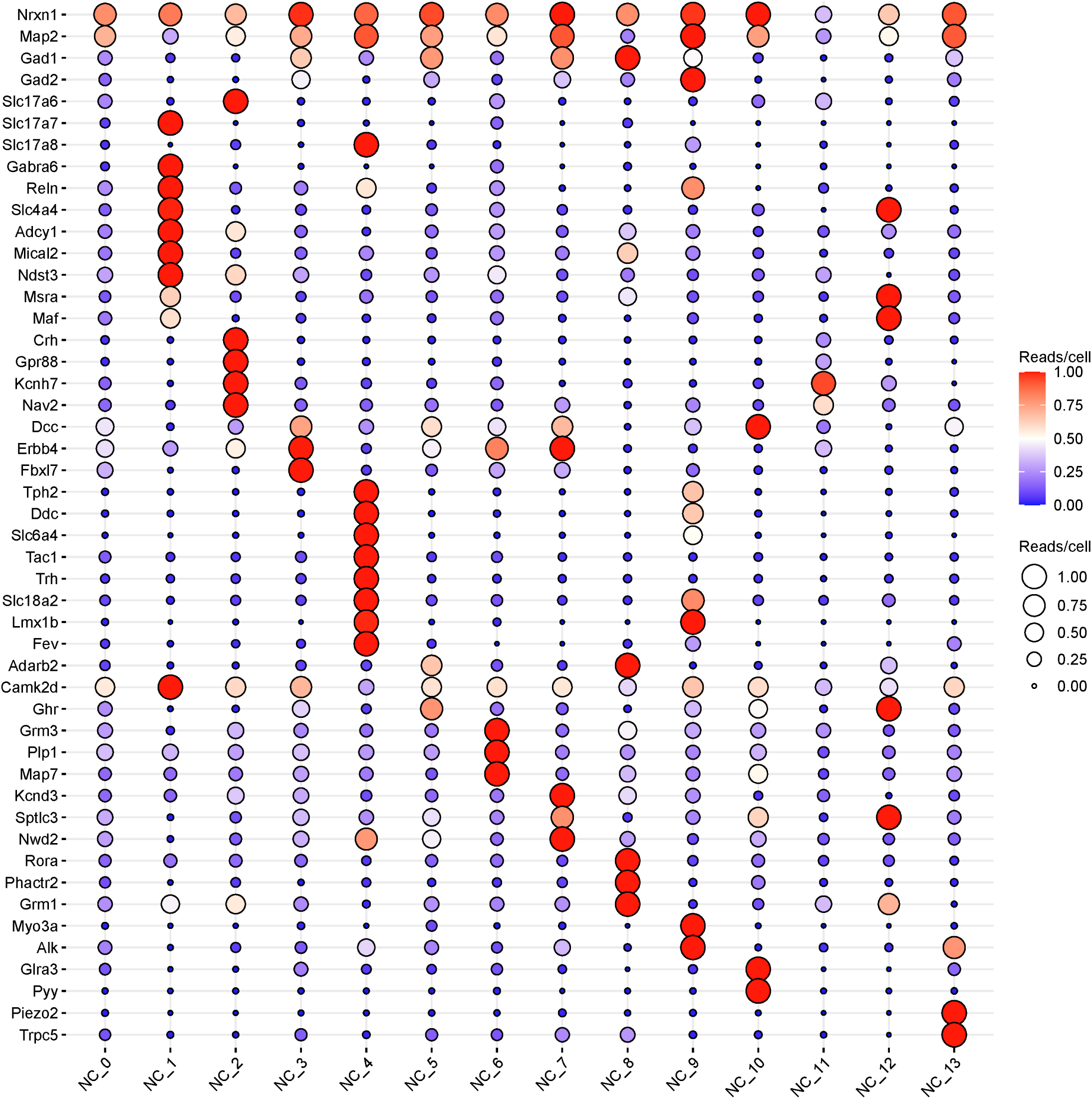
Neuronal Cluster Dot Plot. Expression dot plot of the fourteen distinct neuronal clusters shows unique transcriptional signatures that identify neuronal cell type specificity.

Unique expression of 5-HT neuron genes (*Tph2*/*Ddc*/*Slc6a4*) were found in NC_4 and NC_9, where the largest 5-HT neuron population, NC_4, uniquely co-expressed VGLUT3 (*Slc17a8*) along with *Tac1*, *Trh*, *Slc18a2, Lmx1b* and *Fev* (*Pet-1*; **Fig. 5**). The smaller 5-HT neuron subclass, NC_9, expressed high levels of *Gad2* and co-expressed the marker genes *Myo3a* and *Alk*. This suggested that the majority of the RMg 5-HT neurons captured in this region were glutamatergic and the remaining 5-HT neurons were GABAergic. Additional neuronal clusters were distinguishable by unique combinations of gene markers but were fewer in number. These included: Peptide YY (*Pyy*) neurons (NC_10), which co-expressed *Glra3* and *Dcc*, NC_11 neurons that co-expressed *Kcnh7* and *Nav2*, which was similar to NC_3 but lacked *Erbb4* and *Fbxl7* expression, NC_12 neurons with co-expression of *Slc4a4*, *Msra* and *Maf*, and NC_13 neurons which uniquely co-expressed *Piezo2* and *Trpc5* (**Fig. 5**).

### Distribution of local 5-HT system receptors within the RMg

After identifying subclasses of RMg neurons, we assessed 5-HT system-related gene expression (normalized to maximum expression) across these neuronal cell groups to gain insights into a potential local circuitry. Consistent with previous reports, *Tph2* (not *Tph1*) was expressed in the 5-HT neuron subclasses NC_4 and NC_9, which strongly co-expressed *Ddc, Slc18a2*, *Maoa*, and *Maob* (**Fig. 6**). NC_4 expressed the genes for preprotachykinin (*Tac1*; substance P) and thyrotropin releasing hormone (*Trh*), whereas the receptors for substance P (*Tacr1*; NK-1R) were mainly expressed in local inhibitory neurons NC_5 and NC_7, with relatively less expression in NC_0, NC_2, NC_9 and NC_11. Several 5-HT receptors were expressed in local RMg neuronal subclasses, including a near ubiquitous expression of the 5-HT_2B_ receptor (*Htr2b*). 5-HT_2A_ receptors were expressed highest in NC_13 and less so in glutamatergic NC_2 and the Pyy-expressing NC_10 neurons. Similarly, the other 5-HT receptors were selectively expressed in local RMg neurons, including 5-HT_4_ and 5-HT_7_ receptors (NC_0, 3, 5, 6, 7, 9, 10 and 13), 5-HT_2C_ receptors (NC_7 and 13), and 5-HT_5B_ receptors (NC_2 and 11). However, we found little or no local RMg neuronal expression of other known 5-HT receptors (5-HT_1A/D/F, 3A, 5A, 6_), other tachykinin (*Tacr2*, *Tacr3*) receptors, or the TRH receptor (*Trhr*) in this region. Thus, based on gene expression profiles of 5-HT and/or other receptors across RMg neuronal subclasses, the data provide further evidence of potential local neural circuits that are heavily modulated by 5-HT system neuropeptides.

**Figure 6.**
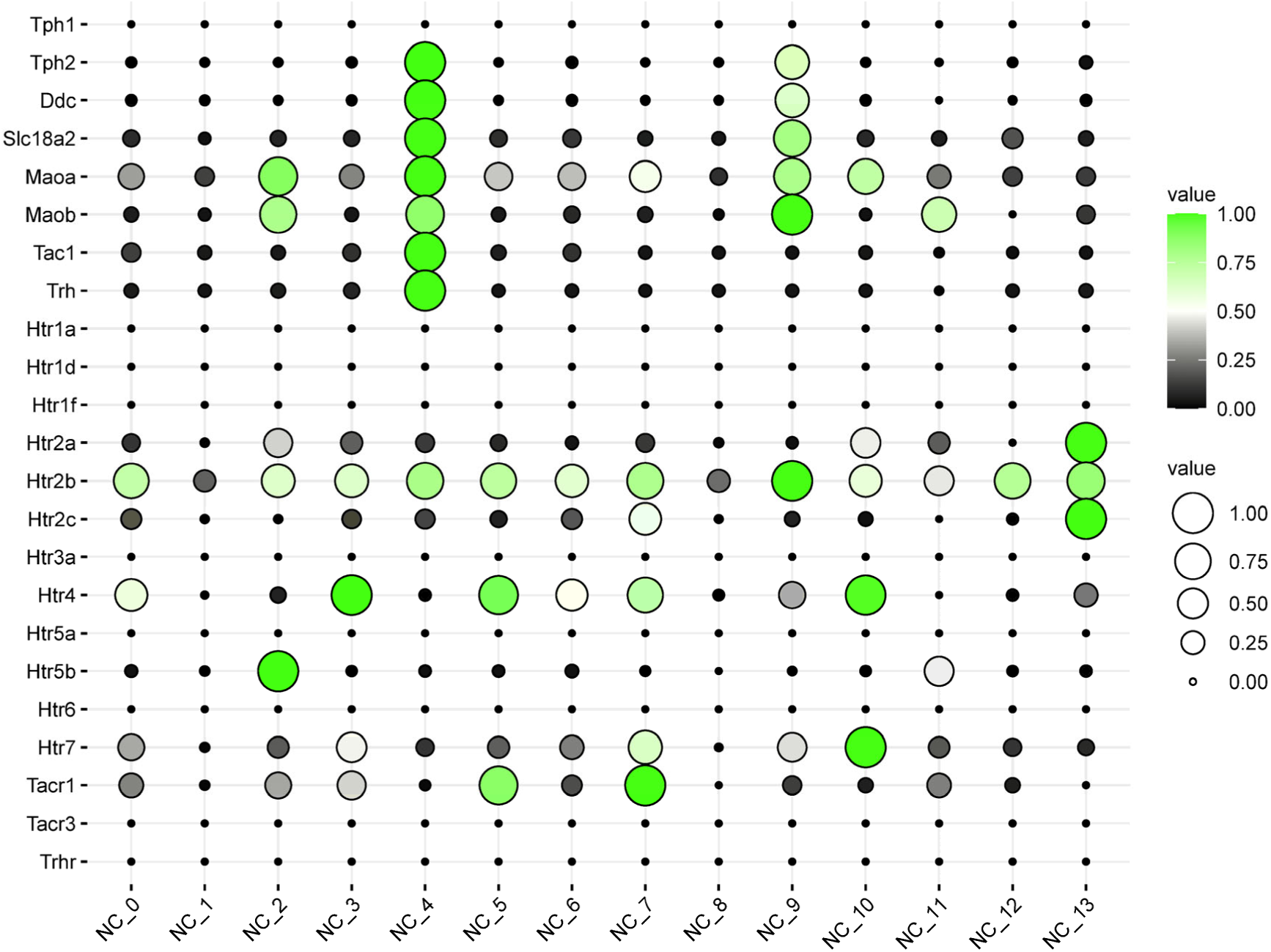
5-HT System Genes across Neuronal Clusters. Cluster 4 exhibits the strongest pattern of 5-HT (serotonergic) neuronal expression in addition to Cluster 9 from the neuron-only reclustering analysis.

### Repeated seizures selectively increase neuronal gene expression

snRNA sequencing has known limitations including sequencing depth, which can limit cell-type specific differential gene expression queries. However, this limitation can be overcome to a degree by combining cell clusters and comparing differentially expressed genes (DEGs) across samples (conditions). We performed two independent differential expression analyses herein to assess any cell type-specific effects of repeated seizures in this model. The first approach combined similar cell types from the original 18 cluster dataset to identify potentially DEGs using standard methods (logFC > 0.5; adj. P value < 0.05; **Fig. 7A**). This approach yielded very few DEGs in non-neuronal cell types comparing tissues collected from rats that had 3, 5, 7, or 10 days of seizures compared to naïve rats (**Fig. 7B**). In contrast, we noted a greater number of DEGs in RMg neurons across all conditions relative to the naïve state. This led to a second approach to identify DEGs in the 14 neuronal clusters (NCs) using the same methodological approach (logFC > 0.5; adj. P value < 0.05), which yielded a very large number of DEGs across all NCs where a majority of DEGs were increased with repeated seizures (**Fig. 8**). These changes in gene expression were essentially ubiquitous across general neuronal cell type and appeared largely independent of how many days of seizures the animals experienced. Thus, it appeared that the large and uniform increase in DEGs within the RMg neurons likely reflected the state of neuronal gene expression at the time of tissue collection (1-1.5h post-seizure) rather than any chronic state of gene expression with repeated seizures.

**Figure 7.**
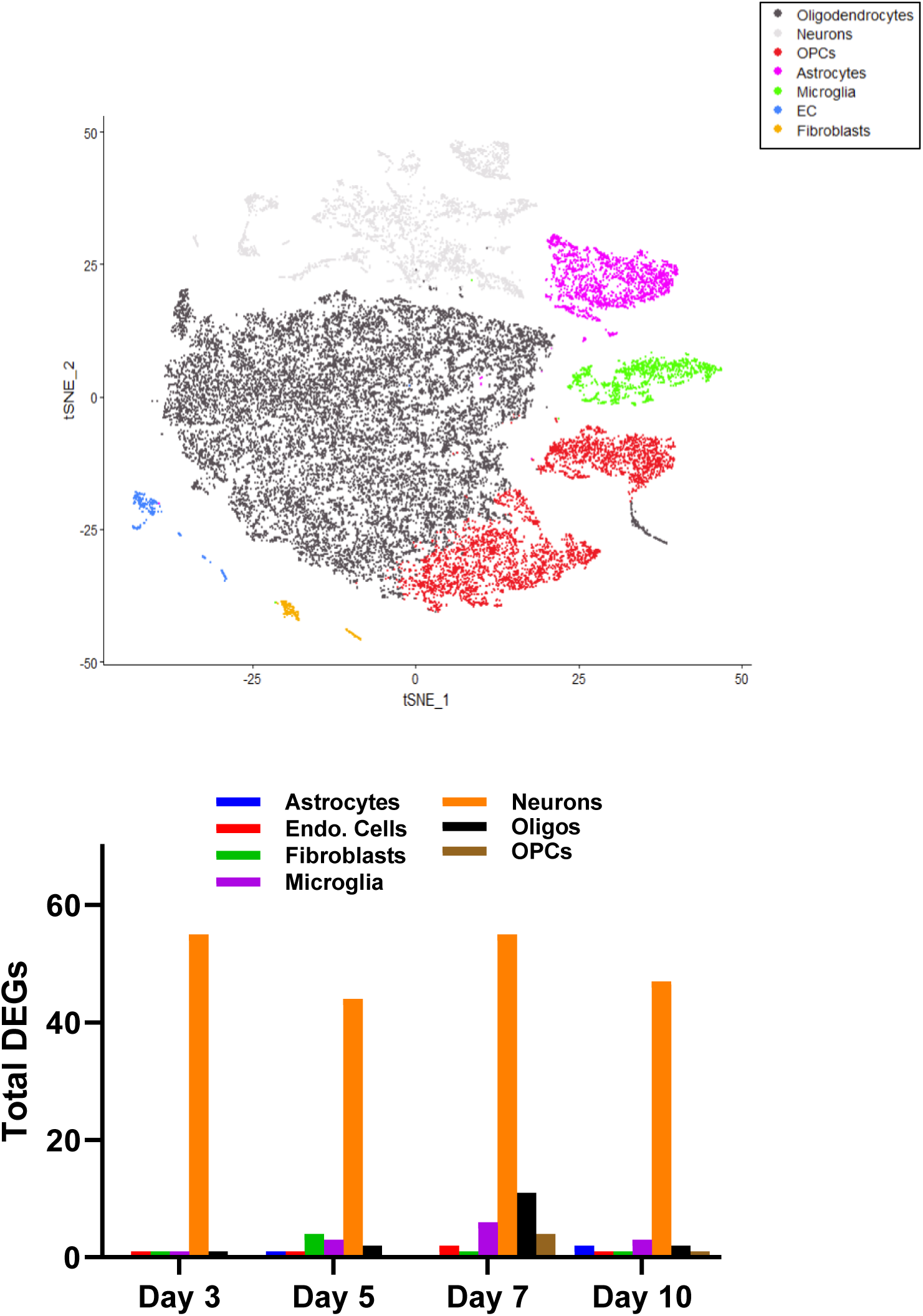
Differentially Expressed Genes Across Seizure Days in All Cell Types. When labeling the t-SNE plot by over-arching cell type, oligodendrocytes predominate. Neuronal cell types consistently had the greatest number of differentially expressed genes after days of repeated seizures with little or no effect in other cell types.

**Figure 8.**
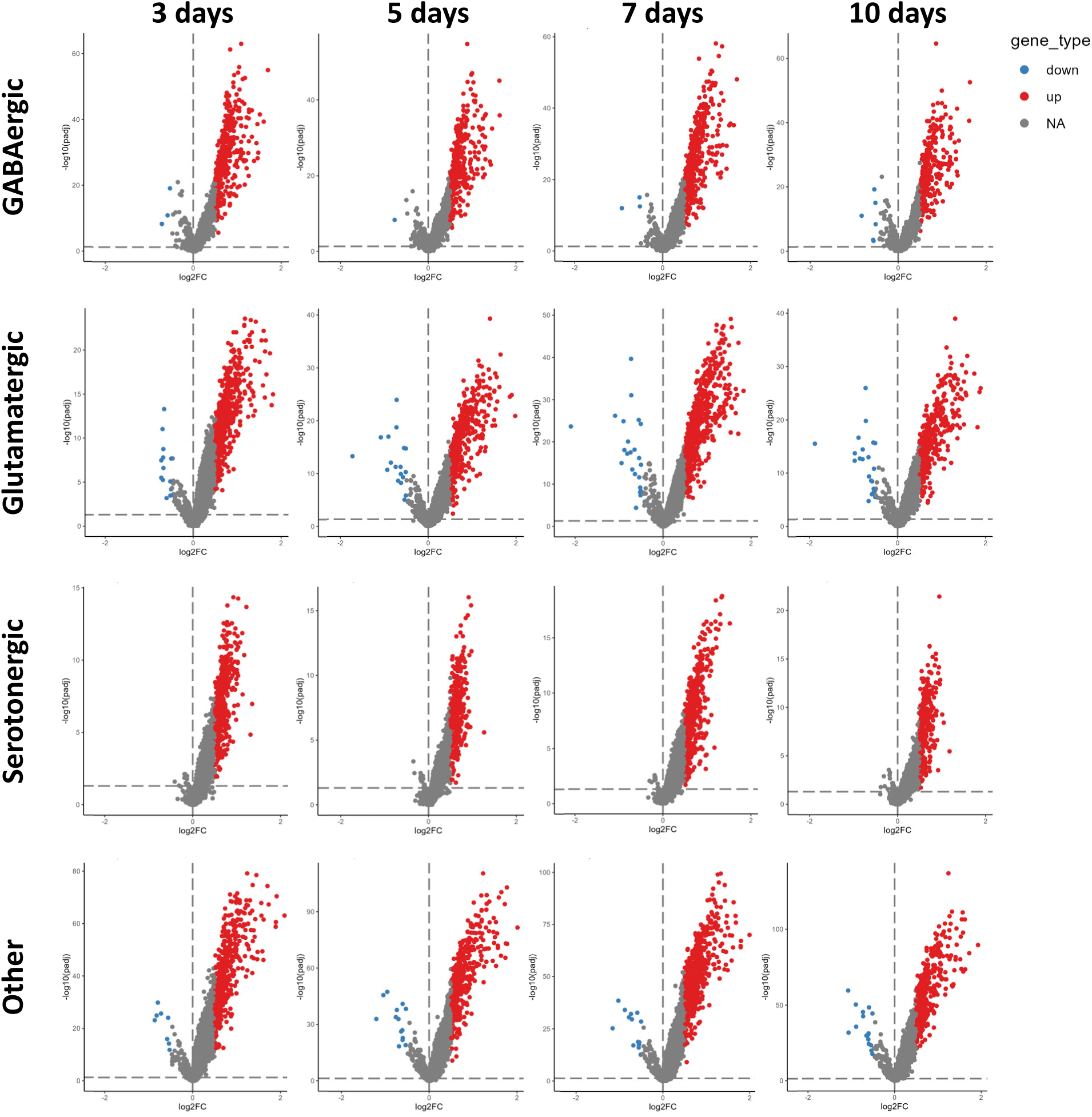
Differentially Expressed Genes in Neurons by Seizure Day. Volcano plots for each of the four neuronal cell types for each of the four seizure conditions. Each neuronal cell-type shows a profound transcriptional shift under all seizure conditions, where the majority of DEGs were increased in expression in neurons after repeated seizures.

The large numbers of genes that increased in expression across neuron type and across conditions appeared very similar in pattern. However, it was unclear how many of the DEGs were in common across neuron type and number of seizures. We mapped DEGs that were common across neuron subtype within days of seizures (**Fig. 9A**) or DEGs common across seizure days within each neuronal subpopulation (**Fig. 9B**). Using this comparison, there was a range of similarity within each condition (21-27%; Fig. 9A) and within each neuron type (23-43%; Fig. 9B), suggesting that there may be a large fraction of DEGs that increased with repeated seizures consistently across seizure days and across different neuronal subpopulations. Using a pathway analysis tool, we next determined which pathways were activated (positive Z-score) and how many DEGs were represented within each pathway (gene count) within each neuronal subtype across all conditions (**Fig. 10**). There were several pathways that reached significance that were shared across all seizure days within each neuronal subpopulation, many of which were related to synaptogenesis which was the top activated pathway in all RMg neuron types (**Fig. 10**). Additional activated pathways in RMg neurons also point to cellular processes involved with activation of cellular remodeling, included Netrin signaling (key mechanism directing cell migration and axon guidance), L1CAM interactions (integral to neurite outgrowth and neuronal adhesion), ROBO SLIT signaling (axon guidance and cell proliferation vs apoptosis), the RHO GTPase cycle pathway (molecular switches directing cytoskeletal remodeling and intracellular signaling), and Axon Guidance signaling. A large fraction of activated pathways also included receptor-mediated intracellular signaling mechanisms, including glutaminergic, GABAergic, opioid, endocannabinoid and NMDA receptor pathways (**Fig. 10**). The presence of these activated pathways across all RMg neuronal subtypes strongly suggest that repeated seizures have several common effects on local neurons, which include but are not limited to cellular processes involved in synaptogenesis, axon guidance, and modulation of excitatory and inhibitory receptors.

**Figure 9.**
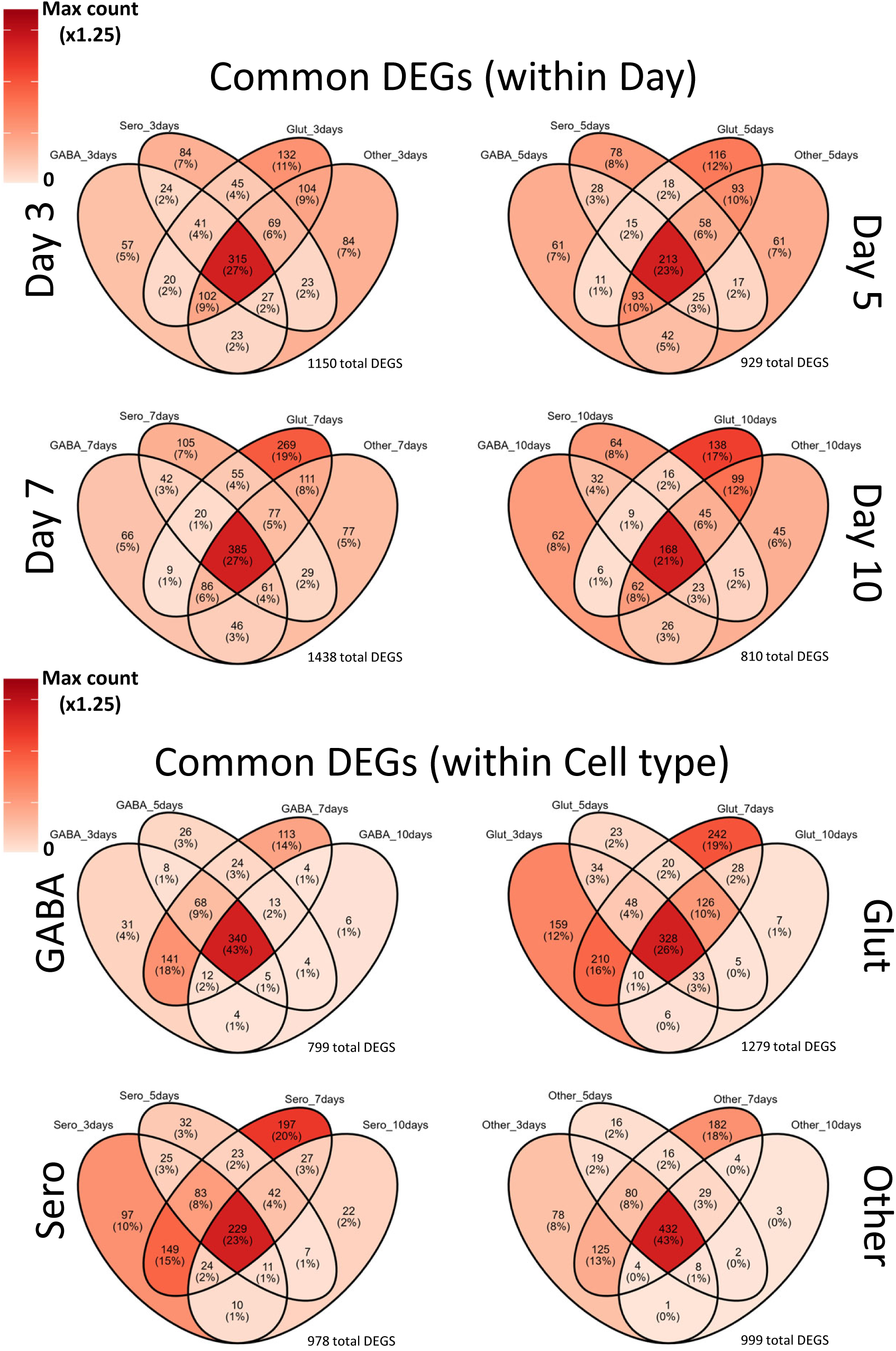
Common Differentially Expressed Genes Across Neuronal Type and Seizure Day. Venn diagrams show that within each discrete seizure day a plurality of differentially expressed genes was shared between the four neuronal cell types, or within each neuronal cell type a plurality of differentially expressed genes that was shared between the four discrete days of seizures.

**Figure 10.** DEGs following repeated seizures drive activation of common pathways across neuronal subtype. Significantly activated pathways (Z-scores>0.5) common to each neuronal subtype across all seizure days highlight synaptogenesis, cytoskeletal remodeling, and modulation of excitatory and inhibitory receptor expression are commonly activated within 1-2 hours post-seizure in the RMg.

## Discussion

The raphe nuclei of the ventral hindbrain, known primarily for containing serotonergic (5-HT) neurons, has been long associated with the regulation of vital systems including breathing, body temperature, and sleep/arousal through 5-HT production^14,20,21^. Brainstem 5-HT neurons arise during prenatal development from transcriptionally-distinct rhombomeres (R1-8), where the large magnocellular 5-HT neurons that populate the Raphe Magnus (RMg) mainly arise from R5^22,23^. These and other hindbrain 5-HT neurons are known to express substance P and thyrotropin-releasing hormone, which together with serotonin provides predominantly excitatory neuromodulatory inputs into cardiorespiratory and thermoregulatory control networks. Deficiencies in aspects of the brainstem 5-HT system including altered 5-HT neuron function or reduced 5-HT levels have been linked to unexpected mortality as seen in Sudden Infant Death Syndrome (SIDS)^24^ and Sudden Unexpected Death in Epilepsy (SUDEP)^6^, suggesting this brainstem region may be critically important in supporting vital functions.

We showed previously that repeated daily generalized tonic-clonic seizures (GTCS) led to a progressive decline in post-ictal ventilation, increased seizure-related mortality and reduced brainstem 5-HT particularly within the RMg^10-12^. However, the complete cellular constituents of the RMg and how these cells may be affected by repeated seizures had yet to be assessed. Here, we identified distinct cell types within the RMg *via* gene expression profiles, including inhibitory GABAergic neurons, excitatory glutamatergic neurons, 5-HT neurons and a small number of Pyy neurons. Further subdivision of RMg neuronal subtypes yielded as many as 14 distinct neuronal subclasses based on combinations of marker gene co-expression, which along with assessment of local 5-HT system receptor genes implicate local neuronal circuits highly modulated by 5-HT neuron neurotransmitters and peptides. Finally, repeated seizures appear to elicit large increases in gene expression in RMg neurons with little or no effects on local non-neuronal cell populations.

The RMg has known roles in several vital control systems, but the functional significance of individual neuronal cell types in each of these physiological functions has been limited in part due to a lack of a complete knowledge of the cellular composition of this region of the brainstem. Unbiased cluster analysis of all cell nuclei collected from the RMg of naïve and seizure-exposed rats provided a first-pass accounting of the cellular constituency in the RMg. Not surprisingly, the 5 transcriptionally-distinct subpopulations of oligodendrocytes represented the greatest number of cells in this region, which could be expected due to the heavy myelination in the brainstem compared to higher brain regions. Neurons were the second most populous cell type, a group comprised of 6 distinct cell clusters of GABAergic, glutamatergic, serotonergic and peptide YY-expressing neurons. Other cell types were identified by known marker genes as oligodendrocyte precursors (OPCs), microglia, astrocytes, endothelial cells, fibroblasts and two additional clusters with mixed transcriptional phenotypes (OPC-Oligo and Fibroblasts_EC). Using snRNAseq provided a complete accounting of the cellular constituency of the rat RMg, thus extending our knowledge of this region of the brainstem.

Reclustering all identified neurons allowed for the identification of 14 distinct neuronal subpopulations by co-expression of unique marker genes. Among the largest populations of neuron subtypes identified within the RMg were two transcriptionally-distinct types of glutamatergic neurons distinguishable by co-expression of either reelin (*Slc17a7*/*Reln*) or corticotropin releasing hormone (*Slc17a6*/*Crh*). While it is unclear what specific functional role these RMg neuronal populations may play, it is important to note that previous data in mice demonstrated CRH expression in 5-HT neurons in the Dorsal Raphe but not in medullary 5-HT neurons^25^ which is consistent with our data. Importantly, neuronal reclustering also yielded two distinct subpopulations of 5-HT neurons distinguishable by the presence or absence of VGLUT3 (*Slc17a8*) and other gene markers of GABAergic neurons. VGLUT3 is known to be expressed in the medullary raphe and most abundantly in RMg neurons involved in thermoregulation^26^. Our RMg 5-HT neuron data are consistent with the concept that some (but not all) monoaminergic neurons express VGLUT3^27^ and may thus influence post-synaptic neurons through fast synaptic (glutamate and GABA) and slower (5-HT, substance P, and TRH) signaling modalities. These synaptic interactions can occur distantly and widely across 5-HT neuron projection fields and/or locally, supported by the data showing 5-HT and NK-1 receptor expression within local RMg neurons. Interestingly, local neuronal NK-1 receptor expression appeared to be largely restricted to inhibitory GABAergic neurons, which appear to modulate important ventilatory chemoreflexes independently from local 5-HT neurons in rats^28,29^. All neuronal subclasses found in this region express one or more 5-HT receptors, strongly implicating functional local neural circuits that are highly modulated by raphe 5-HT neurons through several G protein-coupled receptors^16^.

The VGLUT3-expressing 5-HT neurons identified herein also co-expressed *Tac1* and *Trh,* which were absent in the GABAergic 5-HT neurons suggesting these different RMg 5-HT neuron pools in the rat may arise from distinct embryonic origins (R6P and R5, respectively) as seen in mice^25^. These two 5-HT neuron populations have been shown to modulate ventilatory control mechanisms, particularly as it relates to the maintenance of CO_2_/pH homeostasis. R6P-derived 5-HT neurons co-expressing *Tac1* modulate ventilation during hypercapnia *via* innervation of motor neuron pools for respiratory muscles^30^ whereas R5-derived 5-HT neurons are intrinsically pH/CO2-sensitive^31^ and are GABAergic^25^. We show in our rat data two distinct 5-HT neuron populations with very similar gene expression profiles, suggesting that these neurons also exist in the rat RMg and could functionally contribute to ventilatory control during hypercapnia in rats as they do in mice.

While there we few differentially expressed genes induced by repeated seizures in non-neuronal cells captured in our data, we consistently observed large increases in gene expression in all neuronal subtypes in the RMg. This appeared throughout all RMg neuronal subtypes, which was largely confirmed by comparing common DEGs across days of seizure exposure within neuronal cell type and vice versa (**Figs. 8-9**). Remarkably, these data point to a strong effect of repeated seizures on local RMg neurons with minimal or no effects on other cell types. Furthermore, a large fraction of DEGs across neuronal cell type and numbers of days of seizures were shared, suggesting the potential for a common gene expression pattern in neurons in response to seizures. It is noteworthy that a limitation of this study was that all biopsy samples were collected <90 min post-seizure on the day of collection. This may mean that the gene expression patterns better reflect an acute cellular (neuronal) response to seizures compared to any lasting or chronic transcriptional response. Here we took advantage of this sampling limitation to gain insights into the gene expression state ∼ 90 minutes post-seizure, and found that not only are most DEGs increased in expression level but a major fraction of these genes contribute to the activation of multiple cellular processes that govern synaptogenesis, cytoskeletal remodeling, axon guidance and excitatory and inhibitory receptor expression. Additional future studies are required to better establish potential chronic or steady state changes in gene expression that may result from repeated seizures.

In summary, these data provide a complete cellular constituency of a major brainstem region known for providing excitatory neuromodulatory inputs into vital functions such as cardiorespiratory and thermoregulatory control. These include several neuronal, glial and other cellular subpopulations defined by differential expression of marker genes which may be helpful in designing future, cell-specific manipulations in rats. Furthermore, these data also firmly establish that seizures mainly affect gene expression in local RMg neurons with little or no effects in other cell types in the region, and that there is a large fraction of commonly differentially expressed gene sets across all neuronal subpopulations suggesting a common neuronal response to seizures includes synaptogenesis, cytoskeletal remodeling and shifts in the balance of excitatory and inhibitory receptor expression.

## Supplemental Figures

**Supplemental Figure 1.** Seizure severity scores and the change in body weights of the animals used for snRNA sequencing analysis. **A**) a modified Racine score was used to quantify seizure severity for SSkcnj16-/- rats that did not have seizures elicited (naïve) or those in which 3, 5, 7 or 10 daily seizures were elicited. **B**) Shown are the body weight changes in SSkcnj16-/- rats exposed to repeated daily seizures. There were no differences in seizures severity scores or weight changes across all groups (ANOVA; P>0.05 for days of seizures).

**Supplemental Table 1.** QC metrics of the snRNA sequencing dataset.

