## supplemental figures for "Molecular Deconstruction of the Medullary Raphe Magnus in the rat: Transcriptional Responses to Repeated Seizures"

Suppl. Fig 1. Animal data by group

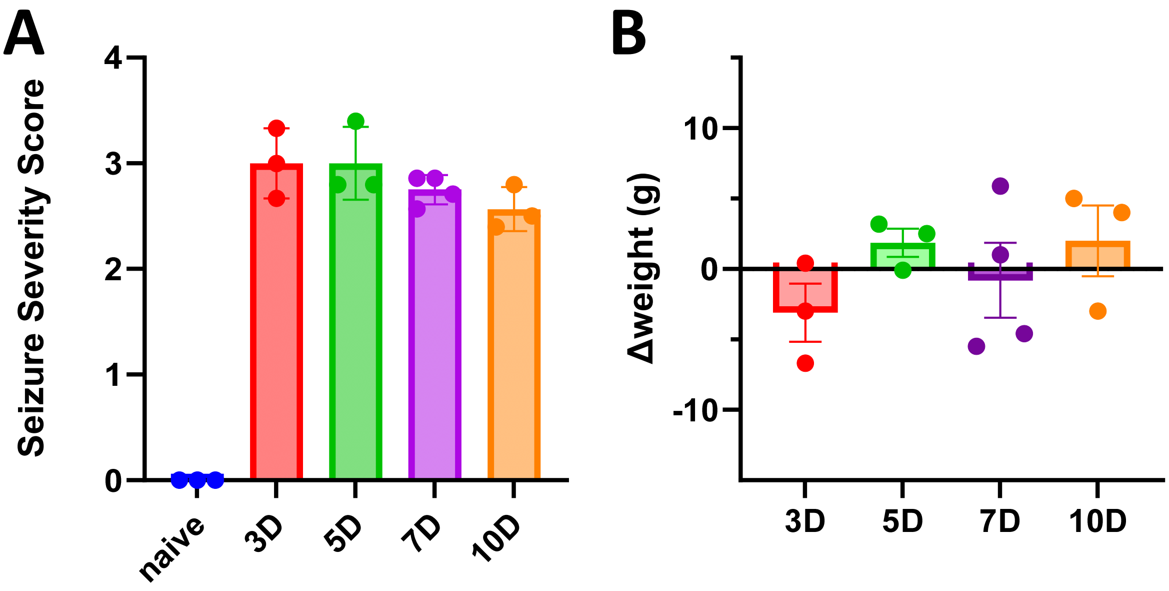

### Suppl. Table 1. QC metrics of single nuclear RNA sequencing

| | **Naïve** | **3D seizures** | **5D seizures** | **7D seizures** | **10D seizures** | **Average** | SD |
|---|---|---|---|---|---|---|---|
| Estimated Number of Cells | 6,573 | 10,661 | 8,469 | 12,836 | 9,118 | **9,531** | 2,358 |
| Mean Reads per Cell | 34,176 | 22,218 | 24,985 | 18,940 | 24,952 | **25,054** | 5,671 |
| Median Genes per Cell | 1,337 | 1,115 | 1,105 | 1,004 | 1,099 | **1,132** | 123 |
| Number of Reads | 224,640,669 | 236,869,473 | 211,595,387 | 243,108,716 | 227,513,232 | **228,745,495** | 12,091,272 |
| Valid Barcodes | 93.6% | 93.8% | 93.8% | 94.0% | 93.8% | **93.8%** | 0.00 |
| Sequencing Saturation | 51.1% | 49.8% | 47.7% | 46.4% | 48.2% | **48.6%** | 0.02 |
| Q30 Bases in Barcode | 95.2% | 95.6% | 95.5% | 95.6% | 95.6% | **95.5%** | 0.00 |
| Q30 Bases in RNA Read | 92.8% | 93.3% | 93.1% | 93.1% | 93.0% | **93.1%** | 0.00 |
| Q30 Bases in UMI | 96.2% | 96.6% | 96.5% | 96.6% | 96.6% | **96.5%** | 0.00 |
| Reads Mapped to Genome | 95.8% | 96.0% | 95.7% | 95.7% | 95.6% | **95.8%** | 0.00 |
| Reads Mapped Confidently to Genome | 88.9% | 88.9% | 88.3% | 88.1% | 88.9% | **88.6%** | 0.00 |
| Reads Mapped Confidently to Intergenic Regions | 10.9% | 10.4% | 11.1% | 10.5% | 11.1% | **10.8%** | 0.00 |
| Reads Mapped Confidently to Intronic Regions | 42.4% | 36.8% | 38.5% | 33.8% | 41.0% | **38.5%** | 0.03 |
| Reads Mapped Confidently to Exonic Regions | 35.6% | 41.8% | 38.7% | 43.8% | 36.7% | **39.3%** | 0.03 |
| Reads Mapped Confidently to Transcriptome | 44.2% | 48.4% | 46.1% | 49.5% | 44.6% | **46.6%** | 0.02 |
| Reads Mapped Antisense to Gene | 33.4% | 29.7% | 30.7% | 27.7% | 32.7% | **30.8%** | 0.02 |
| Fraction Reads in Cells | 43.2% | 48.4% | 41.8% | 46.3% | 42.3% | **44.4%** | 0.03 |
| Total Genes Detected | 21,556.0 | 21,577.0 | 21,501.0 | 21,646.0 | 21,627.0 | **21,581.4** | 57.8 |
| Median UMI Counts per Cell | 1,993.0 | 1,615.0 | 1,555.0 | 1,395.0 | 1,537.0 | **1,619.0** | 224.1 |
